# A comprehensive comparative analysis of bacterial communities within cryoconite holes of Arctic and Antarctic glaciers as a response to global warming changes

**DOI:** 10.1101/2024.09.05.611576

**Authors:** Joanna Sasin-Kurowska, Maciej Golan, Sebastian Piłsyk, Marta Sochacka-Piętal

**Affiliations:** Institute of Genetics and Biotechnology, University of Warsaw, Poland; Military Institute of Medicine, Warsaw, Poland; Institute of Biotechnology and Biophysics, Warsaw, Poland; Rzeszów University of Technology, Rzeszów, Poland

## Abstract

Cryoconite holes, water-filled ponds with sediment at the bottom, play a crucial role for glacial microbiological ecosystems. While they have been widely studied in various regions of the world (e.g., the Arctic, Antarctic, Alps, Himalayas, Central Andes), the novelty of the analyses presented here lies in their deeper exploration of metagenomic data. This study provides a comparative analysis of environmental factors influencing microbial biodiversity, with an additional focus on explaining the effects of pollution on bacterial communities.

## Introduction

Climate warming is causing significant alterations in glacial environments worldwide. Rising global temperatures are profoundly affecting the supraglacial ecosystems, where most of the life in the Antarctic and Arctic regions is microbial. Accelerated glacial melt is expanding microbial habitats within ice sheets and large glaciers, including wet snow, exposed surface ice, hydrologically influenced subglacial environments, and glacial forefields (Cameron et al. 2020). Environmental conditions within these habitats are also altering and will continue to evolve due to rising temperatures, increased rainfall, greater cloud cover, and higher water content (Guo X. et al., 2024; Vavrus et al., 2009). The intensified glacial melt will lead to a greater release of microbiota. Understanding the structure of polar microbial communities and the response of these microbiota to climate change is crucial.

To gain a deeper understanding of the glacial biome, researchers have compared glaciers on both global and local scales. To the best of our knowledge, only a few studies have published comparisons of supraglacial microbial communities. (Cameron et al., 2012; Edwards et al., 2011, 2013; Grzesiak et al., 2015). Here, we present a description of nine different glacial microbiomes and the environmental factors influencing their biodiversity. This paper aims to compare the bacterial communities from two types of polythermal glaciers in Svalbard and Ecology glacier in Antarctica. Hans Glacier is a grounded tidewater glacier that flows into the fjord of Hornsund in southern Spitsbergen, while Werenskiold Glacier, located nearby, is a land-based valley glacier flowing from east to west (Pälli et al., 2003).

This study provides a comparative analysis of how environmental factors affecting bacterial family biodiversity. Additionally, it aims to explain the impact of pollution on bacterial cultures. It extends previous studies on microbiota of these glaciers (Grzesiak et al., 2015, Gawor et al., 2016). The comparative analysis includes three glaciers from the Arctic and Antarctica. Geolocation data from earlier research was used to identify the soil type at each sampling location. Soil type and organic matter enrichment at each site were incorporated into comparative analyses, using organic matter content as a distinguishing factor.

## MATERIALS AND METHODS

### Study areas and sampling

Samples were collected from three model polar glaciers (Ecology-King George Island, Werenskiold - Spitsbergen and Hans Glacier - Spitsbergen). Analyzed data come from three distinct points in the ablation zone– equilibrium line, middle and lower (near the front of glacier). Standard environmental properties, including temperature, a concentration of nutrients (i.e. NO3, NH4, PO4, Si), seston, chlorophyl were measured. Whole DNA was isolated directly from samples of ice. After amplification of the 16S rDNA gene fragments were subjected to sequencing. Sequenced amplicons of up to 1000 nucleotides contained the area of five variable regions V1 to V5. Readings of a poor quality (less than Q20) and readings of less than 700 nucleotides were not considered in further analysis. Sequences were compared by the BLAST algorithm to the reference database of microbial 16S rRNA.

### Statistical analysis

Variation among glaciers in thirty-six environmental factors was investigated through ANOVA, followed by Tukey post-hoc tests. Alpha diversity was assessed using three indices: the Shannon diversity index, which considers both species richness and evenness, the Gini inequality index, commonly used in economics to measure inhomogeneity, and the Margalef Richness Index, that adjusts species richness according to the total abundance per sample. The Gini index ranges from 0 to 1, with lower values indicating a more homogeneous species distribution and values closer to 1 reflecting greater heterogeneity. Mantel tests were conducted to examine correlations. Non-metric multidimensional scaling analysis (NMDS) based on the Bray–Curtis dissimilarity was applied to characterize differences in bacterial community composition between samples (Clarke 1993). We conducted a redundancy analysis (RDA) to quantify the variation in community structures across glaciers and in relation to selected environmental factors. Prior to the analysis, the variables were mean centered within each glacier. All analysis were performed in the R programming environment.

### Identification of pollutions

SourceTracker, based on a Bayesian mixing model (Knights et al. 2011), was employed to identify various sources and estimate their contributions to the bacterial community composition in the cryonites. This tool estimates the relative contributions of microbes from multiple sources to a given environment. In SourceTracker analyses, the relative contributions from different sources to a sink environment are modeled as a probabilistic mixture of source compositions (Baral et al. 2018). Separate SourceTracker analyses were conducted for different taxonomic community levels (total community, abundant community, and rare community) and dominant bacterial phyla/orders.

## RESULTS

The data on 36 environmental factors were collected. ANOVA tests showed that pH (F= 129.56, P < 0.001), calcium hardness (F = 18.7, P < 0.01), SiO2 concentration (F=38.92, p<0.001) in cryoconite holes differed significantly between glaciers. Tukey post-hoc tests showed that glaciers could be divided into two groups with respect to pH values, with Ecology glacier showing higher values than Werenskiold and Hans (|t| >14.5, P ≤ 0.01). With respect to calcium hardness Ecology glacier has much lower values (|t| >8.5, P ≤ 0.01). Similarly, it has also significantly lower concentration values of silicon dioxide (|t| >7.7, P ≤ 0.01).

Bacterial diversity, as estimated by phylotype (order) richness varied across analyzed ecosystems (Table 1). Generalized least squares (GLS) models showed that alpha diversity indices do not differ significantly between glaciers (the values of F statistics p-values are as follows: Margalef Richness Index F=4.23, p>0.05, Shannon index F=1.99, p>0.1, Ginni index F=2.98, p>0.1).

**Table 1.** Characteristic of microbial communities from three polar glaciers.

| Glacier | Distance | Shannon Index | Ginni Index | Margalef Richness Index |
| --- | --- | --- | --- | --- |
| Ecology | 1841.2 | 2.7 | 0.81 | 2.7 |
| Ecology | 457.2 | 2.3 | 0.83 | 2.5 |
| Ecology | 153.8 | 2.24 | 0.84 | 2.52 |
| Hans | 39 | 2.62 | 0.78 | 2.38 |
| Hans | 1760 | 2.1 | 0.83 | 2.3 |
| Hans | 5120 | 2.56 | 0.72 | 1.83 |
| Werenskiold | 50 | 2.09 | 0.85 | 2.18 |
| Werenskiold | 1770 | 2.23 | 0.83 | 2.15 |
| Werenskiold | 3420 | 2.19 | 0.84 | 2.29 |

There is a weak (negative) relationship between Margalef Richness Index and the distance to the glacier’s front (Correlation coefficient equals -0.59, p<0.1). However, the orography of the Hans glacier can have strong impact on results. All the communities were dominated by Actinomycetales, Burkholderiales and Sphingobacteriales bacteria. There are orders specific for particular ecosystems (Xanthomonadales – Ecology glacier, Sphingobacteriales – Werenskiold glacier and Gemmatimonadales – Hans glacier) (Figure 1).

**Figure 1.**
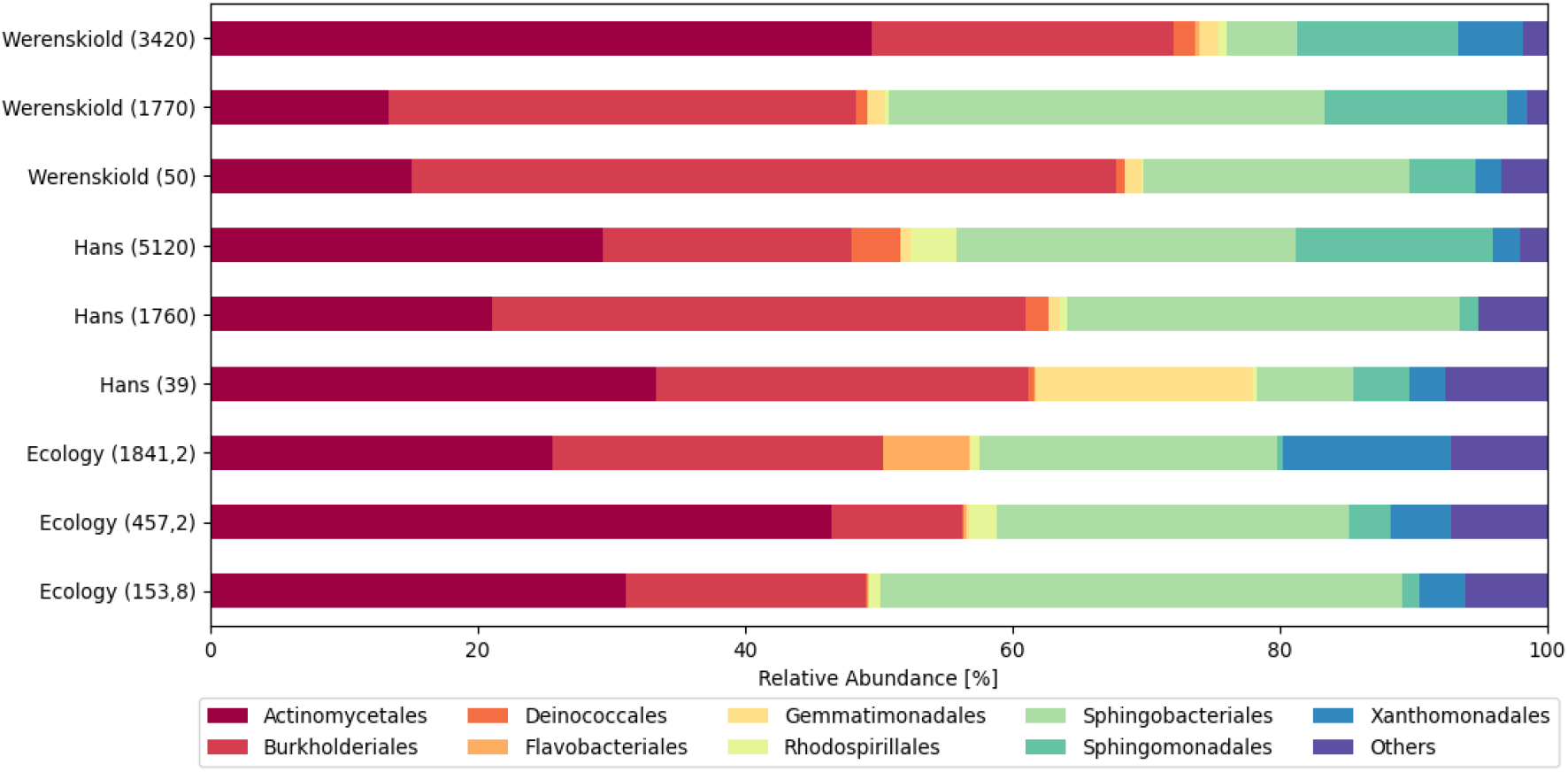
Bacterial composition along the ablation zones of three polar glaciers. Colored bars indicate the percentage of the designated group within each sample.

The SIMPER analysis identified the top 3 orders that cumulatively contributed 72.4% to the differences in bacterial community composition among three habitats (Fig. 2). The top 3 OTUs were dominated by species from orders Burkholderiales, Actinomycetales, Sphingobacteriales.

**Figure 2.**
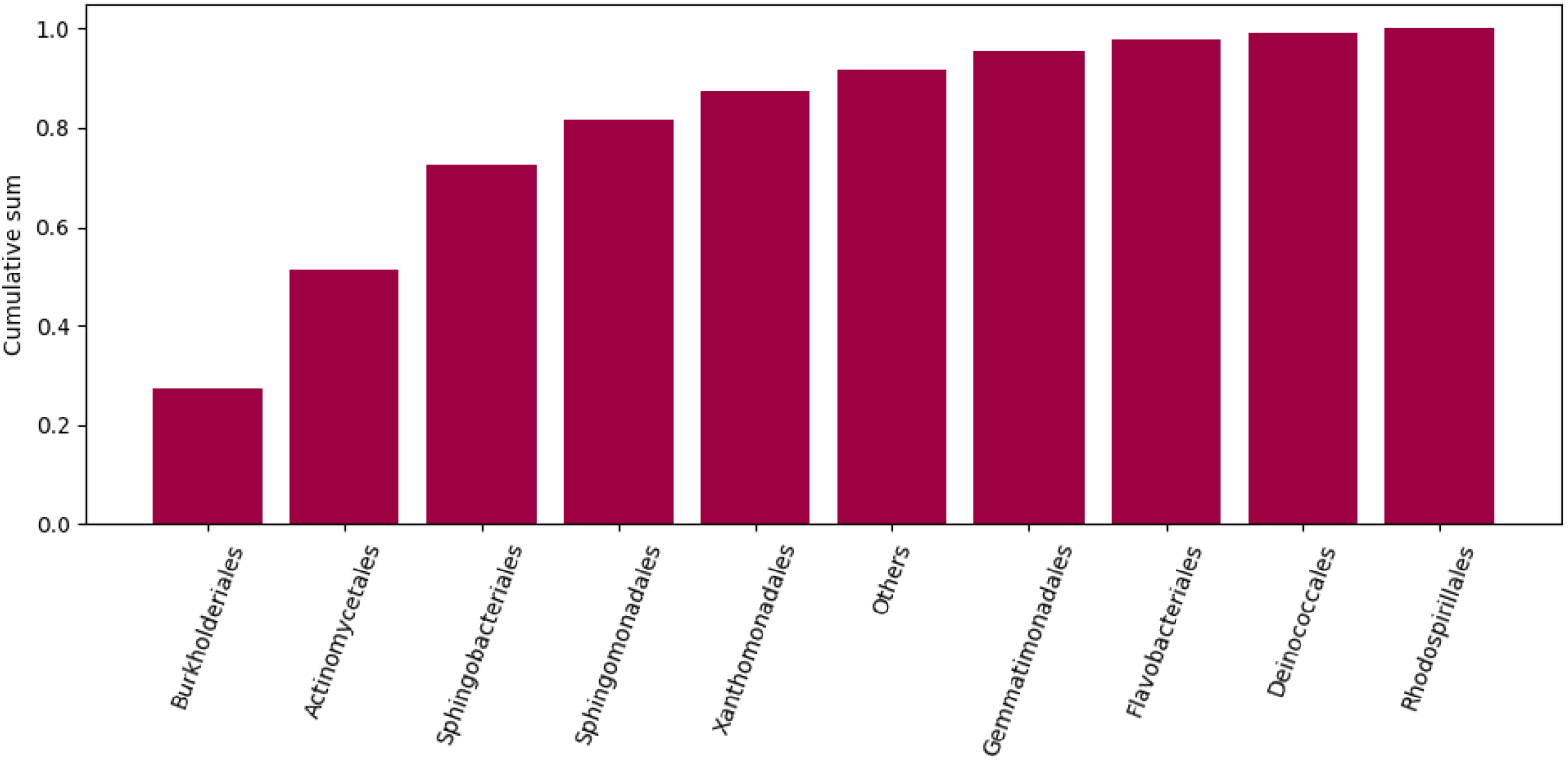
The SIMPER analysis results, represented as a cumulative sum, the principal operational taxonomic units (OTUs) responsible for the differences between habitats belongs to Burkholderiales, Actinomycetales and Sphingobacterials orders.

**Figure 3.**
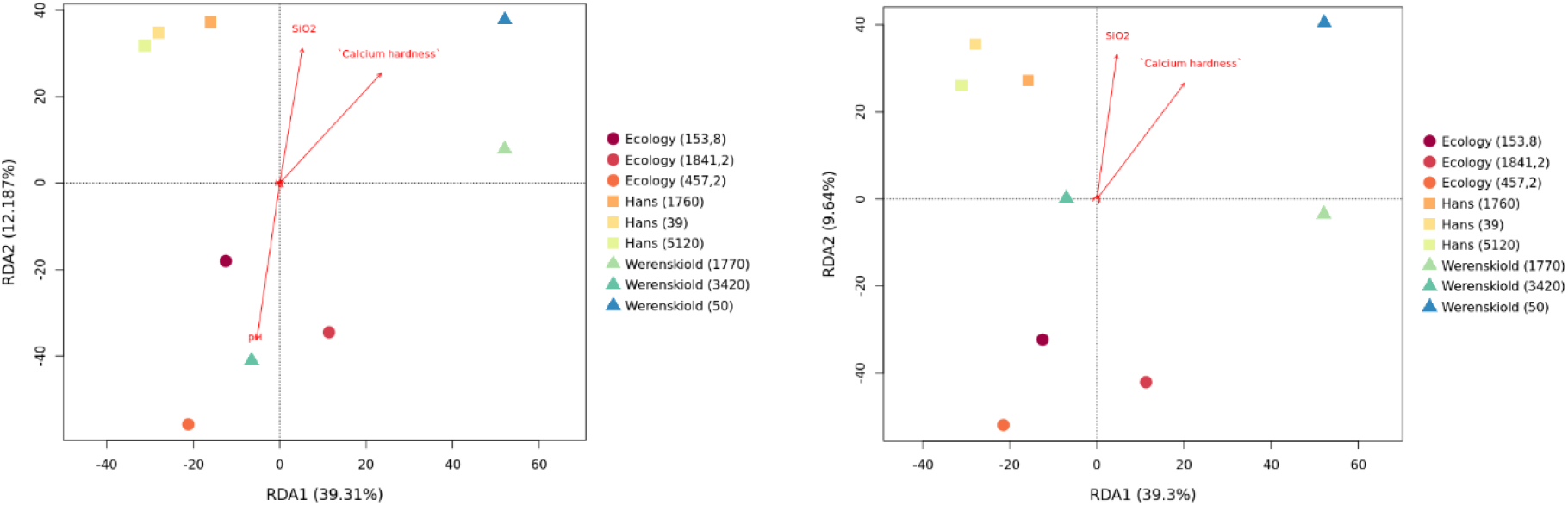
The RDA correlation biplot displays Hellinger-transformed bacterial ASV abundances for all samples, grouped by glacier, pH (only on the left plot), Calcium hardness, and SiO2 concentration, with the latter three or two variables centered to their mean value per glacier. The percentage of variance explained by each axis is reported. The significance of results of the left plot is p<=0.05, while of the right plot is p>0.05, meaning that SiO2, pH and ‘Calcium hardness’ shall be considered as factors that explain variance.

The correlation between the biodiversity indices and thirty-six environmental factors was analyzed. The strongest influence on the composition of microbiota (Shannon index) have the concentration of nitrogen dioxide (r2=0.66, p<0.1), the concentration of sulphate ions (r2=0.62, p<0.1). Another significant environmental factor identified using the Margalef Richness Index is pH (r2=0.78, p<0.05). The concentration of silicon dioxide and calcium hardness have strong negative influence on biodiversity (r2=-0.84, p<0.01, r2=-0,82, p<0.01, respecively).

We conducted redundancy analysis (RDA) to examine the variation in bacterial communities between glaciers and in relation to pH, SiO2, and calcium hardness. This analysis revealed significant differences in bacterial communities among glaciers, with variations associated with pH and SiO2 (Fig. 4). The biplot indicated that bacterial communities in cryoconite holes on the Ecology glacier were primarily influenced by pH, whereas those on the Hans glacier were more affected by SiO2 and calcium hardness. Post-hoc tests further showed that the bacterial community structures across all five glaciers did not differ significantly from one another (F = 2.16, PFDR ≤ 0.058).

**Figure 4.**
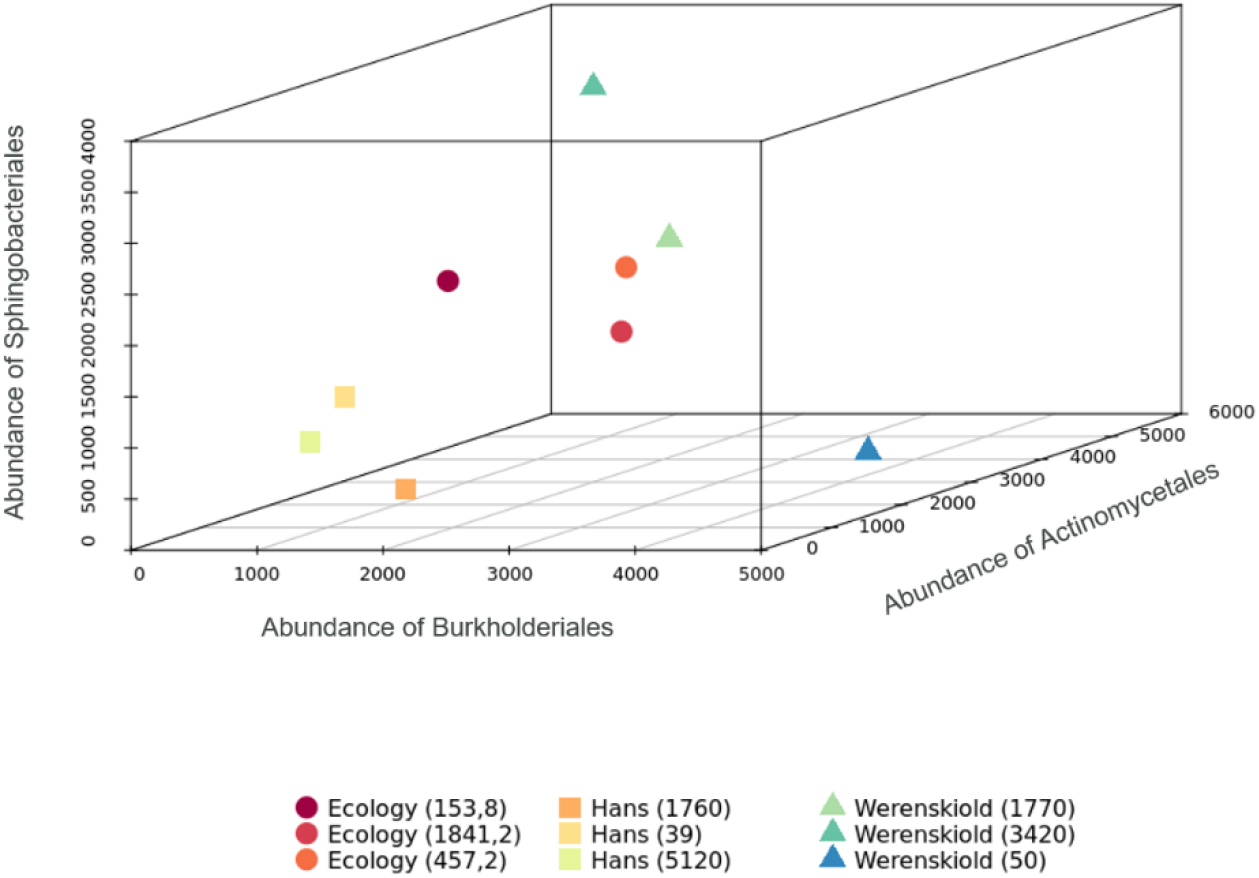
The non-metric multidimensional scaling (NMDS) approach was used to visualize the bacterial community compositions (based on Bray–Curtis distance) across nine habitats. While the abundance of the three principal operational taxonomic units (OTUs) responsible for habitat differences is similar in cryoconites from Ecology and Hans glaciers, it varies in cryoconites from Wersenskiold glacier..

Non-metric multidimensional scaling (NMDS) was used to visualize the bacterial community compositions (based on Bray–Curtis distance) across nine habitats. It revealed the samples from 3 glaciers were clearly separated (Fig. 4). This suggests that the abundance of the three principal operational taxonomic units (OTUs) responsible for habitat differences is more influenced by environmental factors and geographical location than by the distance from the glacier’s front. Therefore, we aimed to examine the impact of pollution on bacterial communities, identify the soil type at each sampling location, and analyze how it influences the results.

## DISCUSSION

This work aims to compare the bacterial communities from two types of polythermal glaciers in Svalbard and Ecology glacier in Antarctica. The results show that from 36 environmental collected factors only pH, calcium hardness, SiO2 differed significantly between glaciers. All the communities were dominated by Actinomycetales, Burkholderiales and Sphingobacteriales bacteria. The SIMPER analysis identified the top 3 orders that cumulatively contributed 72.4% to the differences in bacterial community composition among three habitats (Fig. 2). The top 3 OTUs were dominated by species from orders Burkholderiales, Actinomycetales, Sphingobacteriales. There are orders specific for particular ecosystems (Xanthomonadales – Ecology glacier, Sphingobacteriales – Werenskiold glacier and Gemmatimonadales – Hans glacier).

## Notes

### Competing Interest Statement

The authors have declared no competing interest.

